# Effect of migration and environmental heterogeneity on the maintenance of quantitative variation: a simulation study

**DOI:** 10.1101/226381

**Authors:** Tegan Krista McDonald, Sam Yeaman

## Abstract

The paradox of high genetic variation observed in traits under stabilizing selection is a longstanding problem in evolutionary theory, as mutation rates are 10-100 times too low to explain observed levels of standing genetic variation under classic models of mutation-selection balance. Here, we use individual-based simulations to explore the effect of various types of environmental heterogeneity on the maintenance of genetic variation (V_A_) for a quantitative trait under stabilizing selection. We find that V_A_ is maximized at intermediate migration rates in spatially heterogeneous environments, and that the observed patterns are robust to changes in population size. Spatial environmental heterogeneity increased variation by as much as 10-fold over mutation-selection-balance alone, whereas pure temporal environmental heterogeneity increased variance by only 45% at max. Our results show that some combinations of spatial heterogeneity and migration can maintain considerably more variation than mutation-selection balance, potentially reconciling the discrepancy between theoretical predictions and empirical observations. However, given the narrow regions of parameter space required for this effect, this is unlikely to provide a general explanation for the maintenance of variation. Nonetheless, our results suggest that habitat fragmentation may affect the maintenance of V_A_ and thereby reduce the adaptive capacity of populations.

## Introduction

As genetic variation is the fundamental basis upon which evolution acts, it is important to understand how variation is maintained in order to provide a foundation for answering various questions in biology and related fields, such as missing heritability (Maher, 2008), conservation of biodiversity (Cook & Sgrò, 2017), and population potential to respond to change (Houle, 1992). And yet, the relative importance of factors that influence variation and the mechanism(s) under which it is maintained are not wholly understood (Barton & Turelli, 1989; Mackay *et al*, 2009). The majority of quantitative traits experience stabilizing selection, which in theory should erode genetic variation. However, high levels of standing variation and heritability of quantitative traits are consistently observed in nature (Johnson & Barton, 2005). This paradox – a high degree of genetic variation maintained in the face of stabilizing selection – remains a longstanding, unsolved problem in evolutionary biology and quantitative genetics theory.

The most widely studied explanation for this paradox is mutation-selection balance (henceforth referred to as MSB), the appeal of which lies in its intuitive logic: mutation, as the ultimate source of genetic variation, provides enough input to offset the eroding effect of selection, leading to a state of equilibrium. Under such models, stabilizing selection is assumed according to a Gaussian fitness function with parameter *V_S_* setting the strength of selection on genotypes (where large values result in weaker selection). Multiple MSB models have been proposed, most notably the continuum-of-alleles model from Kimura and Crow (1964) of which two main approximations have been put forth: the Gaussian approximation (Kimura 1965; later expanded by Lande 1976), and the House-of-cards approximation (Turelli, 1984; Bu□rger *et al.*, 1989).

The continuum-of-alleles model makes the basic assumption that for a locus with a continuous distribution of possible alleles, mutation can result in a new allele with an effect different from the pre-existing one. In general, the two approximations differ in the way each handles this assumption. For example, for an arbitrary diploid locus *i* with an allele of effect *x_i_* and a mutation with effect *y_i_*, under the Gaussian approximation the mutated allele would take on a new value which is conditional on the previous state (*x_i_* + *y_i_*). This results in a predicted equilibrium genetic variance 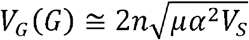, where *n* is the total number of loci that compose the quantitative trait, *μ* the per locus mutation rate, and *α^2^* the variance of the distribution of mutational effects. Under the House-of-Cards approximation, *x_i_* may take on any effect independent from 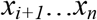, and mutation results in the replacement of *x_i_* by *y_i_*. In this case, the predicted equilibrium variance, 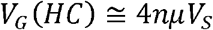 is independent of the distribution of mutation effects. In contrast with the continuum-of-alleles model, there is the diallelic model (Bulmer, 1972; Barton, 1986), which assumes only two possible values at a locus with equal forward and backward mutation rates, such that mutation causes a flip between states (*e.g., xi* = {0,1}). In this case, the resulting equilibrium variance has been shown to be generally comparable to V_G_(HC).

Although these models have been extensively studied, conflicting evidence over their applicability in relation to realistic biologic parameters has left debate open (Barton & Turelli, 1989; Johnson & Barton, 2005; Zhang & Hill, 2005). The assumptions and requirements of these models are unrealistic: primarily, they require extreme mutation rates or too many loci per trait (Johnson & Barton, 2005). This can be most simply illustrated by considering two empirical observations: heritability, *h^2^*, for the majority of traits are relatively high (*h^2^* = 0.2 ~ 0.6; Mousseau and Roff 1987), and stabilizing selection is typically relatively strong in nature, as evidenced by the median value of estimates of reported quadratic selection gradient (γ = -0.1; (Kingsolver *et al.*, 2001). The value γ= -0.1 implies that the ratio of selection to phenotypic variation is V_S_ / V_P_ = 5 (Kingsolver *et al.*, 2001; Johnson & Barton, 2005), which can be rearranged to account for typical estimates of heritability, yielding an expected empirical range of values for variation maintained in a natural population of approximately 0.04 < V_A_ / V_S_ < 0.12. Putting this in the context of Turelli’s (1984) V_G_(HC) model under the assumption that V_G_ ~ V_A_, then 0.01 < *nμ* < 0.03. If there are 100 loci underlying a given trait, then this would require per-locus mutation rates on the order of ~10^−4^, which is 10 to 100 times higher than most estimates from quantitative genetic studies, typically thought to be in the range of 10^−6^ to 10^−5^ (Barton & Turelli, 1989; Bu□rger, 2000).

More complex extensions of these models have been investigated (Bu□rger *et al.*, 1989; Zhang & Hill, 2002; Zhang *et al.*, 2002, 2004), namely by extending to multiple traits, such that ‘apparent stabilizing selection’ is generated through pleiotropic effects. Pure pleiotropy and joint effects models have been studied, as well as other extensions to include factors such as dominance or balancing selection. Nonetheless, these models have yet to provide a sufficient explanation for the patterns of variation maintained as suggested by empirical data (Johnson & Barton, 2005; but see Zhang and Hill 2005).

The above models all assume stabilizing selection to a constant environment, and yet environments varying in space and time can also affect the maintenance of variation. If environmental heterogeneity can maintain sufficient differences in allele frequencies within a subdivided population, and migrant individuals introduce novel variants that can be maintained for some time, then migration can result in an increase in genetic variation within a population. Structured populations with limited amounts of gene flow have the potential to increase within-population variance (Lythgoe, 1997; Tufto, 2000; Spichtig & Kawecki, 2004), and a temporally fluctuating environment has been shown in some cases to increase variance under particular regions of parameter space (Kondrashov & Yampolsky, 1996; Bürger & Gimelfarb, 2002; Gulisija & Kim, 2015). However, these studies have not explicitly framed results in terms of the relative increase in variance over mutation-selection to address the problem of the discrepancy between MSB predictions and empirical observations, and the effect(s) of time and space have not been investigated simultaneously.

Here, we assess how different kinds of environmental heterogeneity affect the maintenance of genetic variation. We simulate spatial and temporal heterogeneity both independently and simultaneously to explore how the maintenance of variation is affected by variation in migration rate, population size, mutation rate, strength of selection, and pattern of environmental variability. In all cases, we explicitly focus on comparing heterogeneous vs. homogeneous environments to represent the increase in variance relative to that expected under mutation-selection balance. This provides some indication of how much more variance can be maintained under heterogeneous environments, which can be compared to the discrepancy between the mutation-selection balance predictions and empirical observations described above.

## Methods

### Simulation Setup

A diploid Wright-Fisher two-population scenario was modeled using the stochastic, individual-based simulation program *Nemo* (*v2.3.45*; Guillaume and Rougemont 2006) under various parameter combinations of population size *N* (1000, 10000), strength of selection *V_S_* (2, 5, 10), mutation rate *μ* (10^−6^, 10^−5^, 10^−4^), and backwards migration *m* (rate ranging from 10^−5^ to 0.1 per generation) between two equal sized patches with some local optima θ. See Table 1 for detailed reference of all parameters.

**Table 1.**
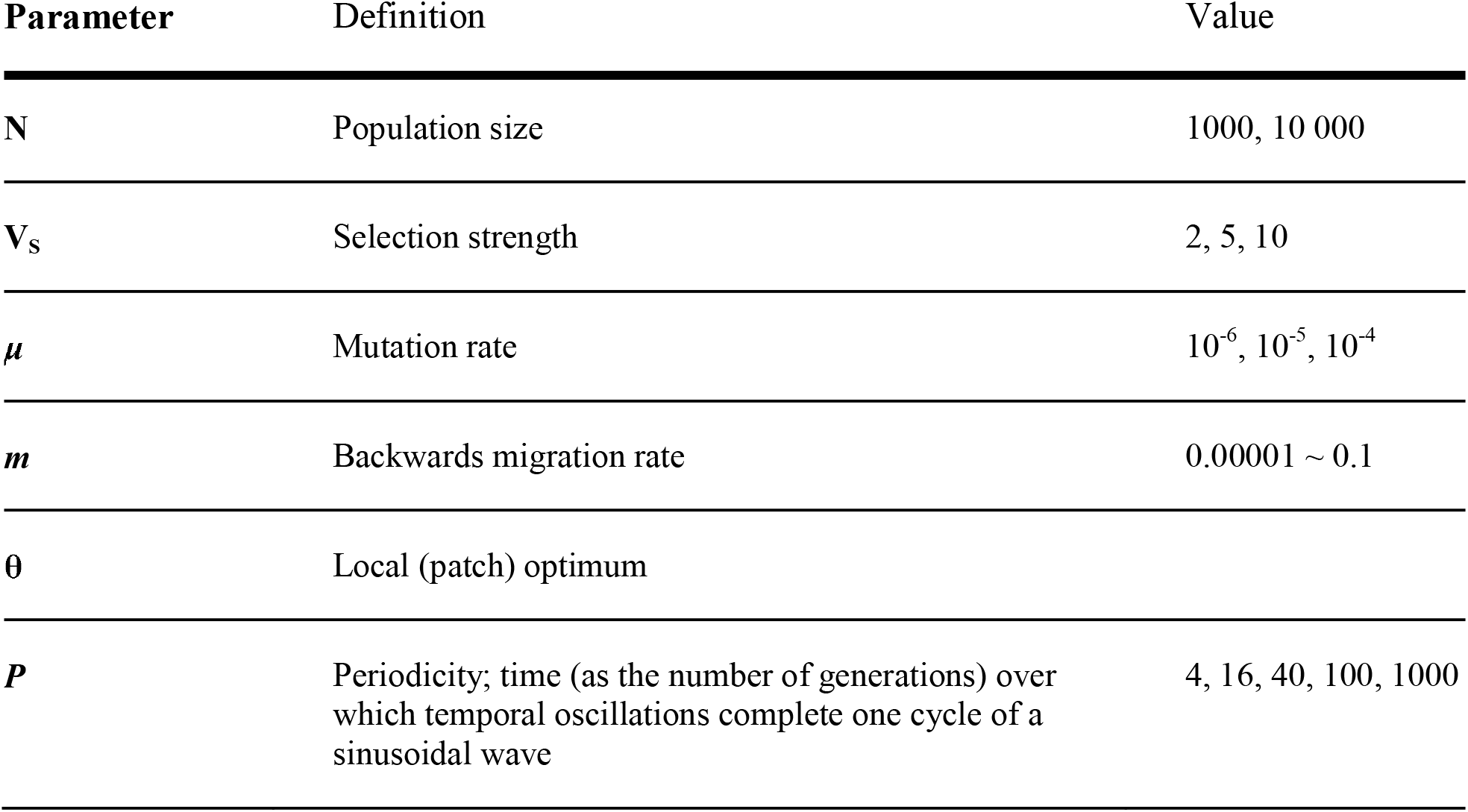
Description of Parameters

In each simulation, relative fitness is determined by Gaussian stabilizing selection acting on a single quantitative trait of 50 unlinked loci, each with an independent allelic value *a* (*n* = 50; *r* = 0.5). Nemo defines individual fitness of a quantitative trait according to the equation:

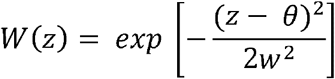

Where *z* is the phenotypic value of the individual, equal to *Σa_i_*, θ the local optimum, and *w^2^* the strength of selection on the phenotype 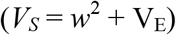. As no effect of environmental variance was included (V_E_ = 0), hereafter we use *V_S_* to represent the strength of selection on the genotype to simplify comparison to theoretical models. The allelic value (*a*) at each locus is randomly drawn from a gamma distribution (mean = 0.05, shape = 1), with mutation causing a change of states ± *a* (this follows a pseudo-diallelic model with a House-of-Cards mutation scheme). Loci have additive effects on the phenotype, with no dominance or epistasis and free recombination between adjacent loci (*r* = 0.5). Thus, genetic architecture cannot evolve through addition of multiple successive mutations at a given locus or through competition among alleles with different linkage relationships.

A total of six scenarios were run, each differing in the relation of local optima between patches:

1. **Homogenous patches** (θ_1_ = θ_2_ = 1). The control set used for comparison to the other scenarios; the local optimum is the same in each patch such that any differences that arise are due to drift rather than selection.
2. **Pure spatial heterogeneity** (θ_1_ = +1; θ_2_ = -1). This set of simulations introduced spatial environmental heterogeneity with opposite local optima between each patch.
3. **Pure temporal heterogeneity** (θ_t,1_ = θ_t,2_). To reflect a population inhabiting a changing environment, temporal fluctuations were modelled using an oscillating optimum defined by a sine wave centered about zero with amplitude of 1; 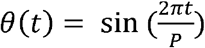. Simulations were run with varying periodicity (P = 4, 16, 40, 100, 1000). The main focus of the analysis considers a temporal oscillation over 100 generations, unless otherwise specified.
4. **Combined (Spatial & Temporal) heterogeneity**. The last set of simulations combined the setup of (2) and (3). Three subsets were used to represent differing degrees of spatial variation (see *SI*): (*A*, θ_t,1_ = -θ_t_,_2_) opposing optima reflected about the horizontal axis; 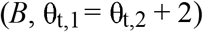 in phase temporal fluctuations with one patch oscillating about -1 and the other about +1; 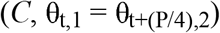 a partial phase shift between optima functions, such that θ_2_ lags behind θ_1_.

All simulations were run for an initial period of 40,000 generations under these conditions, allowing the trait value to stabilize at an approximate equilibrium (assessed visually for the absence of change in trajectory of the trait value over time). A census of the population was taken every *t* generations (t=100, or t=5 for temporal simulations with P ≤ 100 generations) for the mean trait value and genetic variation (in terms of *V_A_*, the additive genetic variation) present in each patch. This data was averaged over the stable period of the last 10,000 generations (1000 for temporal simulations with P ≤ 100) for 40 replicates. For consistency, a 40x50 matrix of allele values was generated such that each replicate had a different complement of mutation effect sizes, but that this complement was held constant across all simulation scenarios. Under the simulations that included temporal heterogeneity, preliminary results showed a cycling pattern emerge for both the trait value and V_A_ over time; to use an estimate as representative as possible, the V_A_ maintained within a cycle was sampled at 20 evenly spaced time-points within a cycle, and the mean calculated over the last ten cycles of the simulation. To obtain a measure of variation maintained within the simulated population that is comparable to empirical evidence, the ratio V_A_ / V_S_ was calculated from the final mean V_A_ estimate of each simulation set. All data processing and analysis was done with R v3.3 (R Development Core Team, 2016).

## Results

### Pure Spatial Homogeneity, Set 1 (θ_1_ = θ_2_ = 1)

We first describe patterns in the homogeneous environment, which acts as a control demonstrating the effect of finite population MSB processes in a two-patch model, to study the effects of mutation rate (μ), selection strength (V_S_), and population size (N), on the maintenance of variation (Figure 1A). As expected from quantitative genetic theory (Falconer and Mackay 1996), variance was somewhat higher in larger populations, although the difference is minimal under most parameters (Figure S1). Also following expectations, V_A_ increases with μ, with the effect of mutation rate approximately linear, and independent of selection. Similarly, V_A_ increases with V_S_ (selection weakens), which qualitatively matches predictions of Turelli (1984), Burger et al. (1989), and as observed by Yeaman and Guillaume (2009), although some deviations occur due to differences in the mutation model. While from Figure 1 there appears to be an interaction between selection and mutation rate, the relative effect of a ten-fold increase in mutation (from μ = 10^−5^ to μ = 10^−4^) is similar for V_S_ = 2 versus V_S_ = 10 (V_A_ changes by a factor of 8.87 vs 8.59 for V_S_ = 2 vs V_s_ = 10; *m* = 0.1).

**Figure 1.**
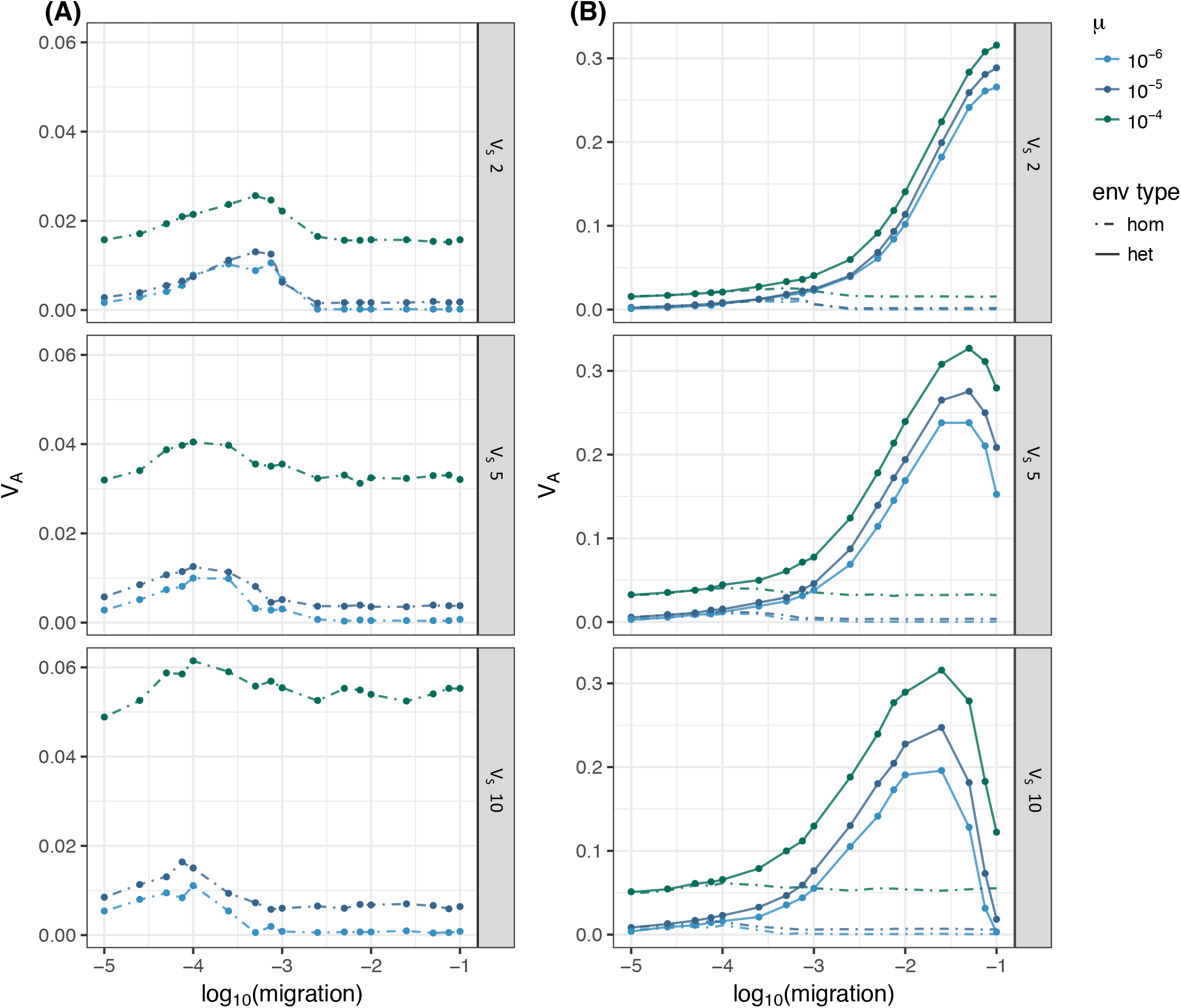
Effect of migration on the maintenance of genetic variation (V_A_) in a sub-divided population (N=1000) between (A) homogeneous patches, (B) spatially heterogeneous patches, under combinations of different mutation rate (line colour; μ={10^−6^, 10^−5^, 10^−4^}) and selection strength (panels; V_S_={2, 5, 10}). The homogeneous environment type (broken line) represents the case of mutation-selection balance.

Notably, these patterns are qualitatively independent of *N*(Figure S1), and so the results below are presented for the subset of N = 1000 only. There is minimal response to rate of migration between the two patches, except for a small increase at low to intermediate *m*, which is consistent with predictions of Goldstein and Holsinger (1992), and arises due to an interaction between effects of genetic redundancy and genetic drift (hereafter, referred to as the “redundancy-drift effect”; see Discussion).

### Pure Spatial Heterogeneity, Set 2 (θ_1_ = +1; θ_2_ = -1)

With spatial environmental heterogeneity, two key differences arise when compared to the control case (Set 1): an overall large increase in the amount of V_A_ maintained, and the appearance of a threshold at high migration, above which the divergence among populations and amount of VA maintained both decrease (which is qualitatively consistent with the critical migration threshold predicted by Yeaman and Otto, 2011). While the migration rate at which V_A_ peaks varies as a function of V_S_, the maximum V_A_ reached under these parameters (≅ 0.32) is similar among levels of V_S_. Given the number of loci, mutation effect sizes, and mutation rates that we used, spatial environmental heterogeneity generates up to a ~10-fold increase in V_A_ compared to the homogenous case (for the same parameters, V_A_[homogeneous] = 0.033), as shown in Figure 1B. This demonstrates that the redundancy-drift effect described previously by Goldstein and Holsinger (1992) is small relative to the effect of migration and spatial heterogeneity, at least under these conditions. Additionally, the effect of mutation becomes relatively less pronounced under spatial heterogeneity, as most of the variation is maintained by migration in this case. Whereas a ten-fold change in μ (from μ = 10^−5^ to μ = 10^−4^) results in an increase by a factor of 8.87 in Figure 1A, this increase only changes V_A_ by a factor of 1.09 under spatial heterogeneity (V_S_ = 2, *m* = 0.1).

### Pure Temporal heterogeneity, Set 3 (θ_t,1_ = θ_t,2_)

In contrast to the case with spatial heterogeneity, under a temporally fluctuating optimum there is in general little change in V_A_ in comparison to Set 1, the homogenous case. As shown in Figure 2A, a similar qualitative pattern is observed in response to migration rate, with an increase at low to intermediate *m* and rapid drop with *m* > 0.001, akin to the redundancy-drift effect observed in the homogeneous case. The resulting maximum peak in V_A_ however is approximately 45% greater than the homogeneous case (V_S_= 5, μ = 10^−4^, *m*= 2.5x10^−4^), suggesting that temporal heterogeneity can only marginally increase the importance of this effect.

**Figure 2.**
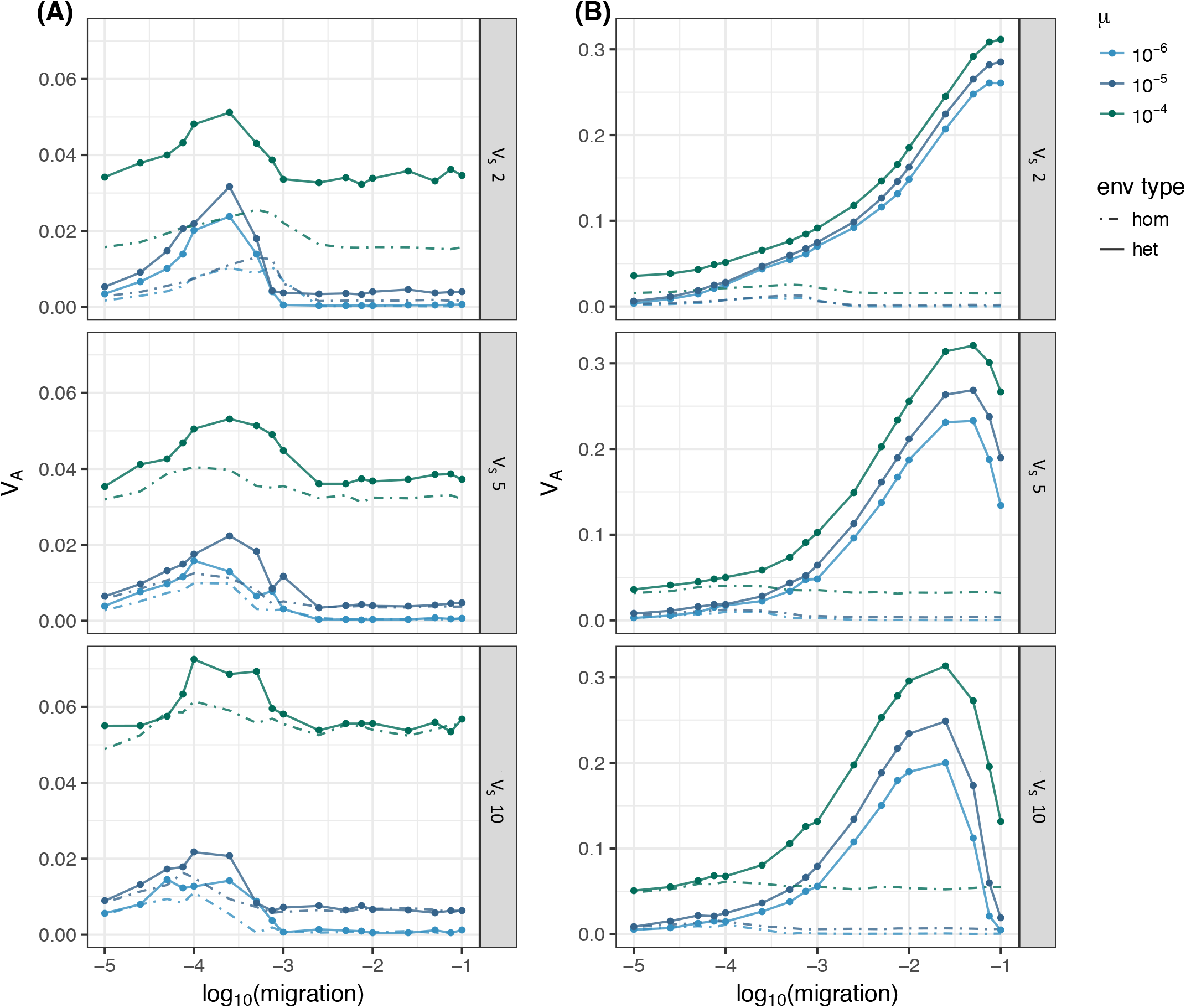
Effect of migration on the maintenance of genetic variation (V_A_) in a sub-divided population (N = 1000) between (A) temporal, (B) combined spatial and temporal, environmentally heterogeneous patches under combinations of different mutation rate (line colour; μ = {10^−6^, 10^−5^, 10^−4^}) and selection strength (panels; V_S_ = {2, 5, 10}). The homogeneous environment type (broken line) represents the case of mutation-selection balance. Period = 100 generations

Multiple lengths for the temporal oscillation of the environment (denoted as P) were simulated to test the influence of the period length on the qualitative patterns observed, and complex interactions were found (Figure S2.1). Setting P ≤ 100 resulted in a response to migration similar to the homogeneous case and an overall increase in V_A_ with μ, as expected. However, the behavior for P = 100 deviates under very strong selection (V_S_ = 2), with the redundancy-drift effect causing a peak in V_A_ at lower migration rates than for shorter periodicities (see Figure 2A and S2.1). Across changes in all parameter combinations, the redundancy-drift effect causes a maximum increase in V_A_ of 28% relative to the V_A_ maintained under *m*= 0.1, which was observed for P = 100 and V_S_ = 10. Under a very long cycle (P = 1000 generations), there is minimal response to migration rate (i.e., no effect of redundancy-drift) and very little increase in variance when μ = 10^−6^ relative to simulations with shorter periods, but much more increase in variance under long periods at μ = 10^−4^ (see Figure S2.1; V_A_ under P = 1000 is 1.3 ~ 2.5-fold higher than P = 100 under μ = 10^−4^). The degree of increase in V_A_ under temporal heterogeneity relative to homogeneity therefore depends on mutation, period, and selection, but is typically considerably less than under spatial heterogeneity with intermediate migration.

### Combination of Spatial and Temporal heterogeneity, Set 4

For spatial and temporal heterogeneity combined (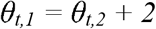; see *SI* for illustration), the amount of V_A_ and patterns that arise are qualitatively similar to those in the case of pure spatial heterogeneity. Similar threshold behavior is observed, and similar levels of V_A_ are maintained, peaking between migration rates of 0.01 ~ 0.1, again depending on V_S_ (Figure 2B). Under this form of combined spatial and temporal heterogeneity, there is a max 31% increase in V_A_ compared to pure spatial heterogeneity (at V_S_= 5, μ= 10^−4^, *m*= 0.001), which is very small when compared to the 9.75-fold difference in V_A_ for the pure spatial heterogeneity vs. homogeneity under the same parameters V_S_ and μ.

In contrast, the complex patterns involving periodicity described above that were observed in the purely temporal set do not appear to carry over under combined spatial and temporal heterogeneity, likely because they are swamped out by the relatively larger effect of migration, which is relatively consistent across different period lengths (see S2.2). There is an overall increase in V_A_ maintained, which appears more similar to the case of pure spatial heterogeneity than pure temporal heterogeneity or homogeneity. This result is perhaps unexpected, as spatial and temporal effects of heterogeneity might be expected to work synergistically.

#### Comparing Sets

Figure 3 shows a comparison amongst the six different patterns of environmental variation for a subset of the total parameter combinations used. For a given parameter set, there is a general trend of increased V_A_ maintained with an increase in heterogeneity, particularly with the degree of spatial heterogeneity between patches. Temporal heterogeneity on its own does little to increase V_A_ compared to the homogeneous set. When temporal heterogeneity is combined with spatial heterogeneity, the increase in V_A_ is most pronounced in sets where the temporal fluctuations in the two patches are most out of phase (Figure 3; see *SI* for relation between θ_t,1_ and θ_t,2_). Of the three combinations investigated, subset 4B – where θ_t,1_ ≠ θ_t,2_ for any *t* — maintains a considerable higher level of V_A_ compared to subset 4A, which in turn maintained more V_A_ than in 4C, which was most in phase between patches. This further supports the emphasis on spatial heterogeneity over temporal heterogeneity as a stronger force for maintaining variation.

**Figure 3.**
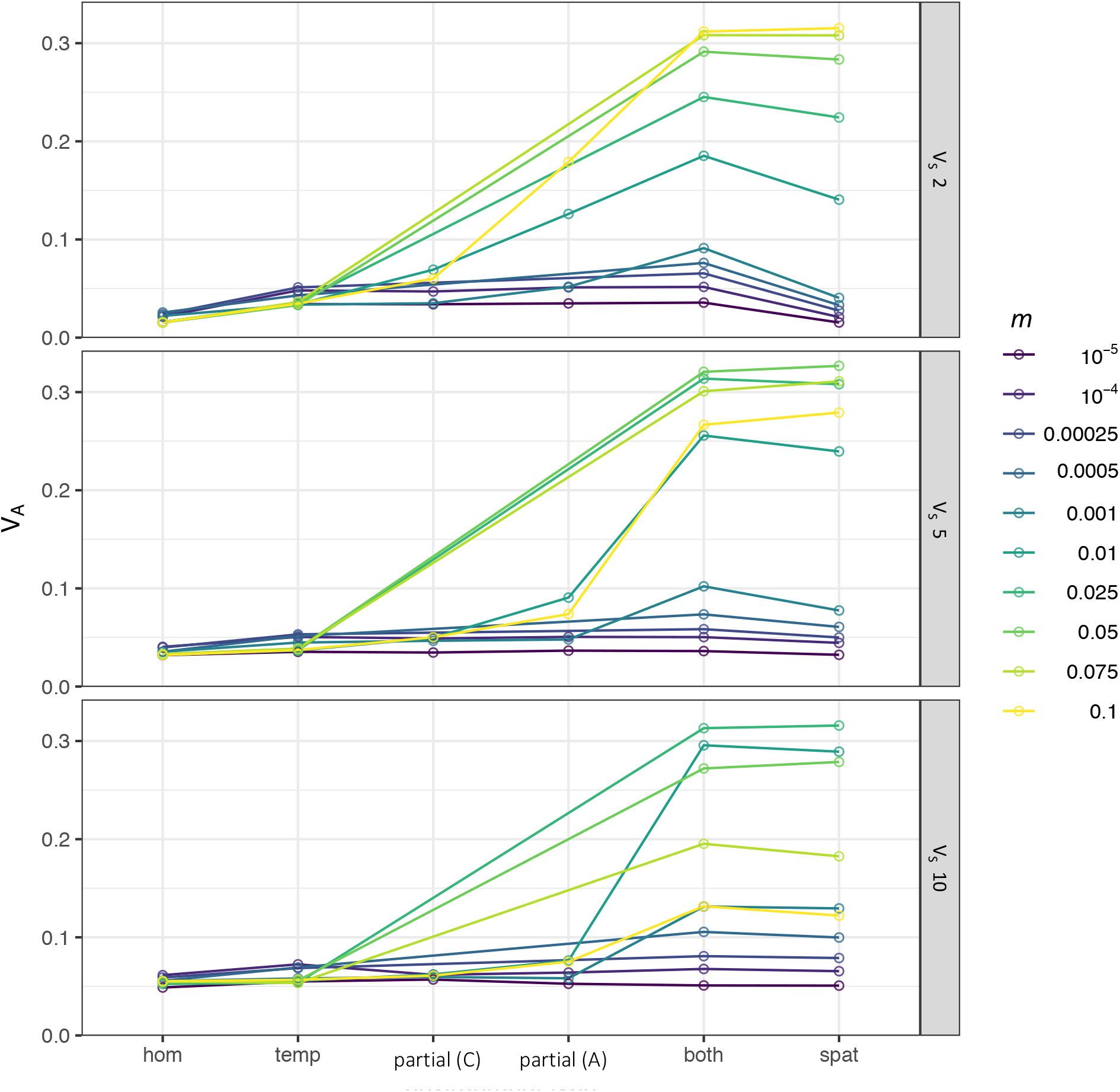
Comparison of the amount of genetic variation (V_A_) maintained in a population simulated under scenarios with increasing degree of environmental heterogeneity, for various rates of migration (*m*) and strength of selection (V_S_ = {2, 5, 10}; panels). N = 1000; μ = 10^−4^, Period = 100 generations

To explore whether environmental heterogeneity could yield more realistic ratios of V_A_ / V_S_ (ie, 0.04 < V_A_ / V_S_ < 0.12), we compared values for this ratio under homogeneous vs. heterogeneous simulation scenarios with all other parameters held constant (Figure 4). Without any spatial heterogeneity, none of the parameter combinations result in V_A_ that falls within this range for expected typical values (see S4, S5 for simulation sets that did not fall into range). However, with spatial heterogeneity, the combination of high migration (*m* ≥ 0.01) and strong selection (V_S_ = 2 or 5) does result in V_A_ within the typical range (i.e. an increase of ~10x relative to MSB). For the combination of spatial and temporal heterogeneity, there is little change for most parameter combinations compared to spatial heterogeneity alone (Figure 4B); for those where there is a noticeable deviation, whether the parameter set falls in or out of the target range remains unchanged (with an exception at V_S_ = 2, *m* = 0.001).

**Figure 4.**
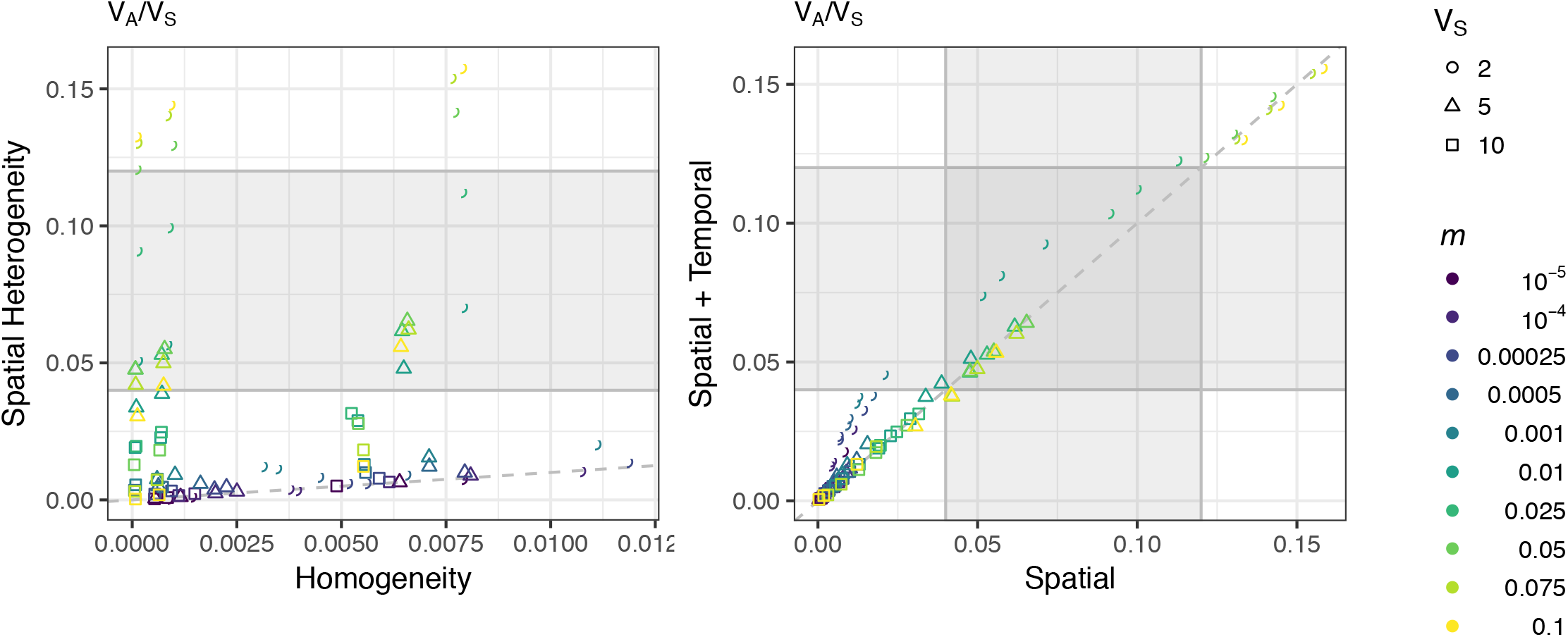
Comparison of the ratio of variance components, V_A_/V_S_, between simulations of different pattern of environmental heterogeneity for various combinations of migration rate (m; point colour), mutation rate (μ = {10^−6^, 10^−5^, 10^−4^}), and selection strength (point shape; V_S_ = {2, 5, 10}). The grey box represents the range of predicted ‘typical’ values expected based on empirical estimates of trait heritability (*h^2^* 0.2 ~ 0.6). Points that fall below the 1:1 line (dashed) are parameter combinations under which more VA is maintained without (A) environmental heterogeneity, or (B) addition of temporal heterogeneity (Period = 100). N = 1000

## Discussion

The results of these simulations show that environmental heterogeneity can have a large impact on the maintenance of additive genetic variation (V_A_) under a variety of parameters. The direction and magnitude of this impact depends upon the rate of migration between subpopulations, with intermediate rates of dispersal providing the largest effect. This effect is also dependent on the type of environmental heterogeneity – spatial, temporal, or a combination of both all result in different magnitude and pattern of response to migration. Spatial heterogeneity has a much larger effect than temporal, and the effect of combined heterogeneity increases with the degree of spatial difference when temporal oscillations in each patch are more out of phase.

### The Redundancy-Drift Effect

For a range of migration rates (dependent on V_S_ and μ) spatial environmental heterogeneity substantially increases the variation a population can maintain. However, migration alone, even between two environmentally similar patches, can increase the variation maintained at very low rates of migration. This result is due to the interaction of genetic redundancy and the stochastic force of genetic drift, as described by Goldstein and Holsinger (1992). With enough population structure (i.e., limited gene flow), the genotypes existing in each patch are expected to undergo different histories of mutation and drift, resulting in relatively independent fluctuations in allele frequency within each patch, and possibly local fixation or loss. Over time, this results in a different pool of genotypes present in each patch, even if the mean phenotype is equivalent. Migration between patches with different allele frequencies will then increase the variance within patches; movement of individuals adapted to a given environment with one complement of alleles that result in a similarly fit phenotype, such that it can be maintained in the new population, then this flux of allelic variants/combinations should increase the variation maintained in the population. This effect is mainly dependent on two factors: (1) the number of possible redundant genotypes, and (2) the degree of population structure – patches must be sufficiently connected such that new genotypes can be introduced by migration at a high enough rate to affect genetic variance, but not overly connected such that drift is no longer acting (relatively) independently in each patch. This genetic redundancy-drift effect is evident in the control case, where there is an observable – albeit minimal – response to low migration, which persists regardless of population size, rate of mutation, or strength of selection. This effect is somewhat increased under temporal heterogeneity, resulting in larger peaks over the same range of low dispersal as the homogeneous case, particularly under high mutation and strong selection (μ = 0.0001, V_S_ = 2). The increase of this effect may be due to the temporally variable selection within each patch causing different alleles to change in frequency more rapidly than migration homogenizes differences, resulting in a larger pool of differentiated loci than would be generated by drift alone.

### Environmental Heterogeneity

Nonetheless, this redundancy-drift effect is small relative to the effect of spatial heterogeneity. In this case, selection acts to pull each subpopulation towards its optima, resulting in different allele frequencies within in each patch. With migration, the alleles that would otherwise be purged by selection are persistently reintroduced into the other patch, increasing the V_A_ maintained in the population. However, there is a threshold behavior to this system – a critical migration rate beyond which polymorphism can no longer be maintained (Felsenstein 1976; Yeaman and Otto 2001). Past this point, the allele frequencies of each patch are similar enough that migrants are much less likely to re-introduce novel/differentiated alleles, and the effect of migration on V_A_ is reduced.

Most populations experience some form of temporally variable selection, be it seasonal fluctuations or some otherwise unstable environment. Due to this, there has been much interest around fluctuating selection and its effect(s) on the maintenance of variation (Kondrashov & Yampolsky, 1996; Bürger & Gimelfarb, 2002; Siepielski *et al.*, 2009; Morrissey & Hadfield, 2012; Gulisija & Kim, 2015). Difficulties arise when attempting to draw comparisons across such studies due to a variety of different assumptions, such as the way in which the moving optimum is modelled and the nature of the underlying genetic architecture. A simple sinusoidal wave is generally used, but varying parameters of this function (amplitude, center, periodicity) appear to have complex interactions with each other as well as other parameters such as mutation rate and genetic redundancy.

Burger and Gimelfarb (2002) found that in a diallelic multi-locus model with recurrent mutation, the relative genetic variance increased with period length (typically ≥ 24 generations), but the magnitude of change and pattern of response to period depended on the amplitude of the oscillations. They observed the highest levels of genetic variance when the amplitude was set such that the optimum cycled between the most extreme possible genotypes, and that greater amplitudes, where the optimum fell beyond the maximum possible genotypic values, resulted in a decline in V_A_ at high period lengths (≥ 52 generations). However, given the small number of loci (2–6) and scaling of allelic effects, this represents a scenario with highly non-redundant genetics. By contrast, Kondrashov and Yampolsky (1996) explored the maintenance of variation under temporal heterogeneity with high genetic redundancy and found that a considerable increase in variance (~3 orders of magnitude) was observed when the amplitude of the fluctuations of the optimum exceeded the width of the fitness function (amplitude,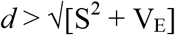). While the assumptions they made about allele effect size and number of loci meant that the local optimum never exceeded the maximum/minimum possible phenotypes, the large amplitude implied by the above inequality results in unnaturally large differences in fitness for a given phenotype over time. For example, an individual with a phenotype equal to +*d* would suffer a fitness cost of 86% when the optimum = *-d*, when *S* = *d* and *V_E_* = 0. Similarly, for the standard set of parameters used in their simulations where substantial variance was maintained, the fitness cost for the same difference in phenotype exceeded 99%. By comparison, the fitness cost for the same difference in phenotype in our simulations was 63% (under strongest selection, V_S_ = 2; this fitness cost is reduced to 18% under weaker selection, V_S_ = 10). Common garden experiments typically show an overall magnitude of local adaptation of ~45% (Hereford 2009), so the selection regimes used by Kondrashov and Yampolsky (1996) constitute much stronger selection than normally observed. Thus, while they found that temporal heterogeneity could maintain much more variance than observed in our simulations, this was likely a consequence of drastic changes in fitness causing the underlying alleles to cycle rapidly in frequency, with increases in V_A_ as the alleles reached intermediate frequency. Thus, the capacity for temporal heterogeneity to maintain substantial amounts of variance seems to depend upon either very extreme environmental fluctuations or a narrow range of rapid fluctuations with strict assumptions on the amount of genetic redundancy available. As empirical data become available for quantitative analyses of a population undergoing temporal variation, this will provide a realistic standard set of parameter values to be used in theory studies, allowing for a more quantitative assessment of the likely effects of temporal fluctuations, and further evaluation of the relative importance of the above models.

#### Spatial v. Temporal v. Combined heterogeneity

Interestingly, it appears that spatial heterogeneity has a much stronger effect than temporal, both in isolation and when combined. This is evidenced by the difference in magnitude of V_A_ maintained in Sets 2 vs. 3 (pure spatial and pure temporal heterogeneity, respectively), as well as the comparisons among the three scenarios of combined spatial and temporal heterogeneity (Set 4 A, B, C). The combination of spatial and temporal heterogeneity only marginally increased the variation maintained by spatial heterogeneity alone, and only in the case of complete spatial heterogeneity (Set 4B). Interestingly, among the three combined sets, V_A_ increased with the spatial component (in general, 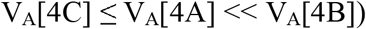), which was robust to changes in μ and V_S_.

Thus, our results indicate that spatial heterogeneity is more important for the maintenance of variation than a temporally fluctuating environment, at least for the genetic architectures and assumptions about fitness used here. Additionally, from the ratio V_A_/V_S_ (Figure 4 and S2), temporal fluctuations alone fail to result in any simulation maintaining significantly more variation than under homogenous conditions, indicating that it is insufficient as an additional mechanism to explain the discrepancy between mutation-selection balance models and empirical observations.

### Does heterogeneity maintain sufficient variation to explain empirical observations?

Given empirical estimates of selection and heritability, a realistic amount of variation maintained in a population should fall within the range V_A_/V_S_ 0.04 ~ 0.12, which would require either an unrealistically large number of loci or very high mutation rates under models of mutation-selection balance, as described in the introduction. To explore whether heterogeneity could result in more realistic ratios of V_A_/V_S_, we compared simulations run under homogeneous vs. heterogeneous conditions, and examined the degree to which heterogeneity increased this ratio. The shift towards this window for simulations with spatial heterogeneity suggests that this is a viable mechanism for the maintenance of variation, but is limited to populations experiencing strong selection and high migration rates.

The results presented in Figure 4 are intended to illustrate how the maintenance of realistic levels of V_A_ is possible under environmental heterogeneity, rather than to represent the full span of parameter combinations that result in realistic behavior. Little is known about the true values for many parameters involved, and changes in some values, such as the number of loci, are likely to alter the quantitative results. Instead, these results illustrate that for a given set of assumptions about the genetic basis of a trait, considerably more variation can be maintained by migration-selection balance than mutation-selection balance. Moreover, the magnitude of this change can be sufficient to explain empirical observations in some cases, indicating environmental heterogeneity is a viable mechanism to consider for the maintenance of variation. However, we stress that given the sensitivity of this result to the balance between selection and migration and the narrowness of the region that maintains significantly more variance, it seems unlikely that environmental heterogeneity will generally resolve the “paradox of variation”.

### Limitations

A major limitation to the interpretation of this study is the difficulty in comparing results to established analytic models due to differences in the mutation scheme. The majority of theory and analytic models assume either a single (or few with equal effect) diallelic loci, or quantitative traits with a normally distributed continuum of alleles. Our simulations attempt to combine the two – varying allelic effects across loci (*a_i_*), but with a single |*a*| per locus, and a Gamma distribution of possible values *a_i_*. We chose this approach to reflect a (potentially) more biologically realistic scheme of mutation. For example, evidence suggests that the distribution of allelic effects at a locus is more likely to follow a gamma rather than a normal distribution, with a small number of loci of large effect and many loci of small effect (Orr, 2003). However, this leads to a less straight-forward parameterization of V_m_ or α^2^, which causes challenges when attempting analytical comparisons.

Another limiting factor is the difficulty of obtaining measurements of migration rates in natural populations, which is necessary to understand if the range where this effect of environmental heterogeneity occurs is realistic. Evidence of migration maintaining quantitative genetic variation in spatially heterogeneous environments was found in lodgepole pine (Yeaman & Jarvis, 2006), but genomic signatures at individual loci may be difficult to detect. Though we did not specifically quantify this here, presumably there is some effect on heterozygosity, though such effects could be difficult to detect in empirical data.

### Implications

We have shown that spatial environmental heterogeneity and migration has the potential to increase genetic variance substantially, while temporal heterogeneity has a much more modest effect. As local adaptation is common (Hereford, 2009), this may be an important factor affecting evolvability. It has been noted that conservation management lacks proper consideration of evolutionary theory, implementation of which can improve overall efforts and long-term outcome (Cook & Sgrò, 2017). The described effect of migration suggests that the ecology underlying the maintenance of variation may be important to consider when planning conservation efforts with a focus on genetics.

For example, a major criterion for the IUCN (International Union for Conservation of Nature) Red List status is population size, however there are examples of listed populations that have remained relatively stable, despite small N_e_ (Ex. Florida Panthers, *Puma concolor coryi;* (Benson *et al*, 2011)). The genetic consequences of population size have been emphasized due to [entirely legitimate] concerns about reduced diversity, which is often used as a proxy measure for the ‘health’ of a population of concern in conservation biology. Here, we showed that a 10-fold change in population size had less effect on V_A_ than a reduction in migration, which suggests that migration may also be an important factor to consider when assessing and managing the genetic health of populations of concern. Indeed, the related concepts of gene flow and connectivity have been previously suggested for consideration and implementation in conservation management, albeit with some contention, as too much migration can result in a reduction of variability (Weeks *et al.*, 2011; Urban *et al*., 2012; Cook & Sgrò, 2017). Long-term maintenance of evolvability may depend in some cases on an intact and interconnected environment, rather than endogenous generation of variation (mutation-selection), which would depend less on environment.

Therefore, a move towards improving connectivity between subpopulations and protection against habitat fragmentation may be a pertinent consideration for the maintenance of quantitative genetic variation, in addition to the well-recognized problems of reducing inbreeding and mitigating demographic fluctuations.

In conclusion, our results demonstrate a clear effect of migration on the maintenance of variation in all four investigated scenarios, and highlight the potential for environmental heterogeneity to substantially increase V_A_. However, it seems unlikely that a single mechanism best explains the maintenance of variation that we see in nature; rather the many concepts put forth over the past decades may be viable explanations in only certain scenarios, just as has been demonstrated here with environmental heterogeneity. Further investigation into potential mechanisms, particularly into those scenarios under which they succeed and fail, and how biologically relevant successful scenarios may be, is needed in order to get a complete picture of the fundamental process that is the maintenance of variation.

## Acknowledgements

We would like to thank D. Lindtke for providing feedback during the writing of this manuscript and M. Turelli for clarifying a theoretical point that SY had muddled. TKM would also like to thank J. Mee for assistance with early stages of the R code. This research was enabled in part by computational support provided by WestGrid and Compute Canada, and was funded by an AIHS grant to SY.

## Literature

Barton, N. & Turelli, M. 1989. Evolutionary Quantitative Genetics: How Little Do We Know. Annu. Rev. Genet. 23: 337–370.

Barton, N.H. 1986. The maintenance of polygenic variation through a balance between mutation and stabilizing selection. Genet. Res. 47: 209–216.

Benson, J.F., Hostetler, J.A., Onorato, D.P., Johnson, W.E., Roelke, M.E., O’Brien, S.J., et al. 2011. Intentional genetic introgression influences survival of adults and subadults in a small, inbred felid population. J. Anim. Ecol. 80: 958–967.

Bulmer, M.G. 1972. The genetic variability of polygenic characters under optimizing selection, mutation and drift. Genet. Res. 19: 17–25.

Bu□rger, R. 2000. The mathematical theory of selection, recombination, and mutation. Wiley.

Bürger, R. & Gimelfarb, A. 2002. Fluctuating environments and the role of mutation in maintaining quantitative genetic variation. Genet. Res. 80: 31–46.

Bu□rger, R., Wagner, G.P. & Stettinger, F. 1989. How Much Heritable Variance Can be Maintained in Finite Populations by Mutation Selection Balance. Evolution (N. Y). 43: 1748–1766.

Cook, C.N. & Sgrò, C.M. 2017. Aligning science and policy to achieve evolutionarily enlightened conservation. Conserv. Biol. 31: 501–512.

Guillaume, F. & Rougemont, J. 2006. Nemo: an evolutionary and population genetics programming framework. Bioinformatics 2556–2557.

Gulisija, D. & Kim, Y. 2015. Emergence of long-term balanced polymorphism under cyclic selection of spatially variable magnitude. Evolution (N. Y). 69: 979–992.

Hereford, J. 2009. A quantitative survey of local adaptation and fitness trade-offs. Am. Nat. 173: 579–88.

Houle, D. 1992. Comparing evolvability and variability of quantitative traits. Genetics 130: 195–204.

Johnson, T. & Barton, N. 2005. Theoretical models of selection and mutation on quantitative traits. Philos. Trans. R. Soc. Lond. B. Biol. Sci. 360: 1411–25.

Kimura, M. 1965. A stochastic model concerning the maintenance of genetic variability in quantitative characters. Proc. Natl. Acad. Sci. U. S. A. 54: 731–736.

Kimura, M. & Crow, J.F. 1964. the Number of Alleles That Can Be Maintained in a Finite Population. Genetics 49: 725–738.

Kingsolver, J.G., Hoekstra, H.E., Hoekstra, J.M.M., Berrigan, D., Vignieri, S.N.N., Hill, C.E.E., et al. 2001. The strength of phenotypic selection in natural populations. Am. Nat. 157: 245–261.

Kondrashov, A.S. & Yampolsky, L.Y. 1996. High genetic variability under the balance between symmetric mutation and fluctuating stabilizing selection. Genet. Res. 68: 157–164.

Lande, R. 1976. The maintenance of genetic variability by mutation in a polygenic character with linked loci. Genet. Res. 26: 221–235.

Lythgoe, K.A. 1997. Consequences of gene flow in spatially structured populations. Genet. Res. 69: 49–60.

Mackay, T.F.C., Stone, E.A. & Ayroles, J.F. 2009. The genetics of quantitative traits: challenges and prospects. Nat. Rev. Genet. 10: 565–577.

Maher, B. 2008. Personal genomes: The case of the missing heritability. Nature 456: 18–21. Nature Publishing Group.

Morrissey, M.B. & Hadfield, J.D. 2012. Directional selection in temporally replicated studies is remarkably consistent. Evolution (N. Y). 66: 435–442.

Mousseau, T.A. & Roff, D.A. 1987. Natural selection and the heritability of fitness components. Heredity (Edinb). 59: 181–197. The Genetical Society of Great Britain.

Orr, H.A. 2003. The distribution of fitness effects among beneficial mutations. Genetics 163: 1519–26.

R Development Core Team. 2016. R: a language and environment for statistical computing. R Foundation for Statistical Computing, Vienna, Austria.

Siepielski, A.M., Dibattista, J.D. & Carlson, S.M. 2009. It’s about time: The temporal dynamics of phenotypic selection in the wild. Ecol. Lett. 12: 1261–1276.

Spichtig, M. & Kawecki, T.J. 2004. The maintenance (or not) of polygenic variation by soft selection in heterogeneous environments. Am. Nat. 164: 70–84. The University of Chicago Press.

Tufto, J. 2000. Quantitative genetic models for the balance between migration and stabilizing selection. Genet. Res. 76: 285–293.

Turelli, M. 1984. Heritable genetic variation via mutation-selection balance: Lerch’s zeta meets the abdominal bristle. Theor. Popul. Biol. 25: 138–193.

Urban, M.C., De Meester, L., Vellend, M., Stoks, R. & Vanoverbeke, J. 2012. A crucial step toward realism: responses to climate change from an evolving metacommunity perspective. Evol. Appl. 5: 154–167. Blackwell Publishing Ltd.

Weeks, A.R., Sgro, C.M., Young, A.G., Frankham, R., Mitchell, N.J., Miller, K.A., et al. 2011. Assessing the benefits and risks of translocations in changing environments: A genetic perspective. Evol. Appl. 4: 709–725. Blackwell Publishing Ltd.

Yeaman, S. & Guillaume, F. 2009. Predicting adaptation under migration load: The role of genetic skew. Evolution (N. Y). 63: 2926–2938.

Yeaman, S. & Jarvis, A. 2006. Regional heterogeneity and gene flow maintain variance in a quantitative trait within populations of lodgepole pine. Proc. R. Soc. B Biol. Sci. 273: 1587–1593.

Yeaman, S. & Otto, S.P. 2011. Establishment and maintenance of adaptive genetic divergence under migration, selection, and drift. Evolution (N. Y). 65: 2123–2129.

Zhang, X.S. & Hill, W.G. 2005. Genetic variability under mutation selection balance. Trends Ecol. Evol. 20: 468–470.

Zhang, X.S. & Hill, W.G. 2002. Joint effects of pleiotropic selection and stabilizing selection on the maintenance of quantitative genetic variation at mutation-selection balance. Genetics 162: 459–471.

Zhang, X.S., Wang, J. & Hill, W.G. 2004. Influence of Dominance, Leptokurtosis and Pleiotropy of Deleterious Mutations on Quantitative Genetic Variation at Mutation-Selection Balance. Genetics 166: 597–610.

Zhang, X.S., Wang, J. & Hill, W.G. 2002. Pleiotropic model of maintenance of quantitative genetic variation at mutation-selection balance. Genetics 161: 419–433.

